# The whole genome analysis of four Orf virus strains from Europe and South America

**DOI:** 10.1101/2024.09.25.613117

**Authors:** Marco Cacciabue, Javier Moleres, Laura C. Lozano Calderón, Irache Echeverría, Lorena de Pablo, Idoia Glaria, Guido König, Andrea Peralta, Ramsés Reina

## Abstract

Orf virus (ORFV) is the etiological agent of Contagious Ecthyma (CE), a worldwide disease that mainly affects sheep, goats, wild ruminants, and humans. Here, we determined the complete genome sequence of two ORFV strains from Spain (NAV and ARA) and two from Argentina (HRE and CHB), representing the second report from Europe and the first from South America. The assembled genomes of the ARA, CHB, HRE, and NAV strains of ORFV were found to be 137,891 bp, 137,160 bp, 137,340 bp, and 137,214 bp long, respectively, containing 132 genes in each strain, showing high amino acid identity and similar lengths to the reference strain NZ2. We performed a microsatellite analysis to determine a molecular signature for strains that differ in their host (sheep or goat). In addition, the analysis of 32 specially selected genes showed that the nucleotide substitution rate for this dataset was 3.2x10-5 subs/site/year (95% HPD 4.2X10-8 -7.6 X10-5) with a median value of 2.8 x 10-5 subs/site/year, placing the TMRCA (median divergence time) in 1661. The genetic characterization of ORFV strains not only allows epidemiological studies but is the first step toward the development of molecular tools oriented to diagnostics and vaccines.

## Introduction

Contagious Ecthyma (CE), also known as Orf, contagious pustular stomatitis or contagious pustular dermatitis, is a worldwide-distributed viral skin disease affecting primarily goats and sheep and wild ruminants but also humans causing self-limiting painful pustular lesions on fingers and hands (McElroy et al. 2007). CE is caused by Orf virus (ORFV), a member of the family *Poxviridae*, and the more prevalent within the Parapoxvirus (PPV) genus, which also includes bovine papular stomatitis virus (BPSV), pseudocowpoxvirus (PCPV), PPV of red deer in New Zeland (PVNZ) and PPV of the grey seal (Mercer et al. 1999). ORFV infects epithelial cells causing severe proliferative dermatitis, which evolves from macules, papules, pustules to scabs and fissured crusts (Nandi S et al 2011). Lesions are commonly found around the lips, mouth muzzle, nostrils, teats and oral mucosa, but can be also found in esophagus, hooves, reproductive orgns, rumen or respiratory tract (Zhao et al. 2010). Albeit lesions get usually resolved in 1 to 2 months, reinfection is commonly observed (McKeever DJ et al 1988), regardless the administration of live-attenuated vaccines.

ORFV infection is endemic in most countries with sheep/goat-rising industries (Robinson, Balassu 1981). Affected animals, mainly young lambs, dramatically reduce their food intake due to painful lesions occurrence, leading to transient impairment in daily weight gain, and therefore, to important economic losses in farms with recurrent outbreaks. Noteworthy, morbidity rate often reaches 100%, and despite mortality rate is usually low, secondary bacterial infections, i.e., pododermatitis and mastitis (Burriel, 1997), further increase sanitary costs (Haig and Mercer, 1998)

ORFV genome consists of 132 putative genes distributed along 135kbp linear double-stranded DNA (Delhon et al., 2004) with an unusually high GC content (∼64%) under positive selection (Sahu et al., 2020). Recently, recombination has been suggested to be the main force driving genetic evolution and ORFV virus diversity (Sahu et al., 2020), jeopardizing vaccination strategies. Strikingly, despite worldwide distribution and increasing Orf outbreak reports, few strains have been isolated, a small proportion of them have been completely sequenced and little genetic information of the related strains is available. There are more than 20 whole genome sequences publicly available, from which we have selected 19, including isolates from China, North America, Germany, France, India and New Zealand (see **table 1** for strains and references).

**Table 1.** Strains of ORFV, PCPV and BSPV used in this study.

| Strain | Country of isolation | Year of isolation | Host | Accession number | Reference |
| --- | --- | --- | --- | --- | --- |
| HN3 | China | 2012 | Sheep | KY053526 | Chen *et al* 2017 |
| NA1 | China | 2015 | Sheep | KF234407 | Li *et al* 2015 |
| GO | China | 2018 | Goat | KP010354 | Chi *et al* 2005 |
| YX | China | 2015 | Goat | KP010353 | Chi *et al* 2005 |
| SJ1 | China | 2015 | Goat | KP010356 | Chi *et al* 2005 |
| NP | China | 2015 | Goat | KP010355 | Chi *et al* 2005 |
| SA00 | USA | 2004 | Goat | AY386264 | Hosamani *et al* 2009 |
| IA82 | USA | 2004 | Sheep | AY386263 | Hosamani *et al* 2009 |
| NZ2 | New Zealand | 2006 | Sheep | DQ184476 | Mercer *et al* 2006 |
| D1701 | Germany | 2011 | Sheep | HM133903 | McGuire *et al* 2012 |
| CL18 | China | 2018 | Sheep | MN648219.1 | Not published |
| NA17 | China | 2016 | Goat | MG674916 | Zhong *et al* 2018 |
| TVL | USA | 2019 | Sheep | MN454854 | Heare *et al* 2020 |
| SY17 | China | 2016 | Sheep | MG712417 | Zhong *et al* 2018 |
| MP | India | 2017 | Goat | MT332357 | Sahu *et al* 2020 |
| VR634 | - | 1963 | Human (Cow) | GQ329670.1 | Hautaniemi *et al* 2010 |
| F00.120R | Finland | 2000 | Reindeer | GQ329669.1 | Hautaniemi *et al* 2010 |
| HRE | Argentina | 2018 | Sheep | ON805831 | This study |
| CHB | Argentina | 2018 | Sheep | ON805830 | This study |
| NAV | Spain | 2018 | Sheep | ON805832 | This study |
| ARA | Spain | 2018 | Sheep | ON805833 | This study |

Phylogenetic studies usually rely on singles genes, being *orf011* (B2L) followed by *orf020* (VIR), *orf132* (VEGF) and *orf059* (F1L) the most reported in the literature (Chan et al., 2007; Guo et al., 2004; Hosamani et al., 2006). According to VGEF alleles (*orf132*), a classification in NZ2 and NZ7-like strains was proposed (Mercer et al., 2006). More recently, according to *orf011* and *orf020* genes, a classification attending to animal species, sheep or goat has been proposed (Karki et al., 2019; Velazquez-Salinas et al., 2018). However, depending on the selection pressures affecting different genes, evolution inferences may suffer from inaccuracy when considering individual or concatenated genes (Yao et al., 2020).

Microsatellites, also known as simple sequence repeats (SSR), are 1-6pb unit repeated ubiquitously in the genome of eukaryotes, prokaryotes and viruses (Alam et al., 2013; Davis et al., 1999; Mrázek et al., 2007; Tóth et al., 2000). Compound microsatellites (cSSR) are composed of two or more individual SSRs directly adjacent to each other, for example (CAG)n-Xn-(TA)n. Some studies have proposed the use of SSR and cSSR for the characterization of viral strains (Sahu et al., 2020; Walker et al., 2001).

In this study, we have analyzed four different CE outbreaks reported in South America and Europe. ORFV presence was confirmed in all animals by performing specific PCR amplification of *orf045* and *orf011* genes in lesions. Isolation was attempted in different ovine primary cells using scabs and replication assessed by molecular methods. Complete genome sequences were obtained by NGS methods and analyzed regarding genome structure, SSRs, recombination events, evolutionary rate and phylogenetic inferences.

Genetic characterization of ORFV circulating strains may shed light on the molecular epidemiology that underlies ORFV infection. Deciphering genome content and diversity is a first step towards development of new detection and prevention tools against EC.

## Material and Methods

### Sheep flocks and tissue collection

Spanish flocks from Navarra (42° 49′ 00″ N 1° 39′ 00″ W) and Aragón (41° 39′ 23″ N 0° 52′ 36″ W) under routine veterinary surveillance, and after reporting two outbreaks of Contagious Ecthyma in April and May 2017 respectively, were included in the study (sample NAV and ARA, respectively).

The Argentine samples were collected in Huinca Renancó (34 ° 50′22 ″ S 64 ° 22′19 ″ W) province of Córdoba (sample HRE) and Chacabuco (34 ° 38’31’’S 60 ° 28’17 ’’ W) province of Buenos Aires (sample CHB) in 2015 and 2016, respectively. In both cases, shepherds did not report previous cases of Contagious Ecthyma, with the difference that CHB flock incorporates new animals every year, while HRE is from a closed establishment.

Scabs were excised from animals by veterinarians using scalped blades and tweezers and maintained refrigerated until shipment to the respective laboratory and then at - 80°C for further analyses.

### Virus purification and DNA extraction

Scabs were thoroughly homogenized in liquid nitrogen and 20 mg of the resulting tissue powder were placed in a microcentrifuge tube and incubated with lysis buffer (100mM Tris-HCl pH7.5, 12.5mM EDTA, 150mM NaCl, 0,5% SDS) and proteinase K at 56°C in a water bath, until complete tissue lysis. DNA was then extracted, according to the manufacturer’s instructions (EZNA DNA tissue kit) and stored at −20°C.

For ORFV detection, PCRs covering 045 and 011 genes were carried out using previously described methods (Kottaridi et al., 2006).

### Sample preparation for MiSeq sequencing

In order to purify viral particles from the scabs, a previously developed protocol (Zwartouw et al., 1962) was used with some modifications: For Argentina samples, five hundred milligrams (500mg) of scab material were macerated in a pestle under liquid nitrogen until a homogeneous powder was obtained. A 30% suspension (weight/volume) was made in TMN buffer (10mM Tris-HCl pH 7.5; 1.5mM MgCl_2_; 10mM NaCl). In the case of Spanish samples, one-half of the scab macerate (50mg) was resuspended in MEM-D medium and used as inoculum to infect ovine epithelial primary cell cultures (Eov). Two passages in Eov cells were made and cells and supernatants from both passages were harvested, and added to the second half of the crust sample in TMN buffer. Samples were freeze-thawed at -80°C once and sonicated three cycles of two minutes each, in bath (Elmasonic, sweep mode). Then, the suspension was centrifuged at 2000 x g for 10 min at 4°C. Clarified supernatant was loaded onto a 30% sucrose cushion (30% w/w sucrose in TMN buffer) and pelleted at 39000g for 30min (rotor 70Ti Beckman). Pellet was resuspended in 2ml of TMN buffer. Additionally, this virus preparation was further purified in an increasing sucrose cushion (30-60%) by centrifuging at 12° C in swinging rotor SW41 at 39,000g for 20min. Purified virus formed an opalescent halo in the 50% sucrose layer. The purified virus was collected and diluted with 2 Vol of TMN buffer and centrifuged at 35,000g for 60min. The pellet was resuspended in 50μl of TMN buffer and stored at - 80°C.

### Genomic DNA preparation and sequencing

DNA extraction was performed according the manufacturer’s instructions of QIAamp DNA mini Kit (QIAGEN). DNASeq library compatible with short read Illumina sequencing was generated using the NEB Ultra DNA library Kit (NEB) starting with 500 ng of DNA, as measured by Qubit (Invitrogen) and following the manufacturer’s instructions. Briefly, DNA was fragmented, end-repaired and subsequently the adapter was ligated. Agencourt AMPure XP beads were used to size select the DNA fragments containing the adapters. Finally, the library was amplified by 15 PCR cycles. The fragment size distribution of the library was analyzed on a BioAnalyzer High Sensitivity LabChip showing a size range between 400 and 446 bp with the main peak of the library at 401 bp. The library was diluted to 2 nM and multiplex-sequenced together with five samples on the Illumina MiSeq (2 × 250 bp paired end run, estimated 4.3 million reads/sample).

### Raw sequence data processing, mapping, assembly, and genome annotations

For each sample, the sequenced raw data were processed to obtain high-quality reads. Reads with a quality score of q30 or lower as well as those under 50 bp were discarded and adapter-trimmed using BBDuk (Bushnell, 2019). Sequences were mapped against the host genome (*Ovis aries*) to remove host DNA contamination. Unmapped reads were used as input data for *de novo* assembly using SPAdes genome assembler v3.15.2 (Bankevich et al., 2012). This procedure rendered two (NAV) or three (HRE, CHB and ARA) contigs that corresponded to ORFV virus according to Blastx searches. The resulting contigs were aligned to the reference genome NZ2 (DQ184476.1) and manually checked to obtain draft genomes. Finally, each set of high-quality reads was aligned to the corresponding draft genome using Bowtie2. The consensus genome sequences were extracted using bcftools (Danecek et al., 2021).

New ORFV whole sequenced genomes were annotated using GATU with NZ2 as genome reference to capture all the potential open reading frames (ORFs) (Tcherepanov et al., 2006).

### Detection of Simple Sequence Repeats (SSRs)

Simple and compound microsatellites were extracted with IMEx software (Mudunuri et al., 2010), which identified perfect mono-, di-, tri-, tetra-, penta- and hexanucleotide repeats along the genomes characterized. The minimum numbers of iterations were 6, 3, 3, 3, 3 and 3 for mono- to hexanucleotide motifs, respectively, using the parameters previously used for RNA (Chen et al., 2009) and DNA viruses (Hatcher et al., 2015; Wu et al., 2014). Maximum distance allowed between any two SSRs (dMAX) was 10 nucleotides. Other parameters were used as default. Compound microsatellites (cSSR) were not standardized in order to determine real composition.

### Phylogenetic analysis

The whole genome sequences obtained were aligned using the MAFFT version 7 package (Katoh and Standley, 2013) and curated manually. The phylogenetic analysis was carried out through the maximum likelihood method using IQ-Tree program version 1.4.2(Minh et al., 2020). IQ-Tree was also used for estimating the substitution model by means of ModelFinder. Then, the tree was built using GTR+G4 evolution model within 1000 replicates for bootstrap, and the results were visualized with FigTree 1.4.0 tool (available at https://github.com/rambaut/figtree/releases)

Alternatively, MrBayes version 3.2.7 (Ronquist et al., 2012) was used for building trees through Bayesian methodology, setting nst: 6, rates: gamma, and invariant sites as model parameters. This analysis was run for 10 million Markov chain Monte Carlo (MCMC) iterations, sampling trees every 1000 generations. 10% the burn-in was considered. Tracer 1.7 version (Rambaut et al., 2018) was used for checking the convergence and evaluating that the effective sample size for relevant parameters was higher than 200. Finally, the tree topology was visualized with FigTree.

Sequence clusters were determined using the Fast hierarchical Bayesian analysis of population structure algorithm (fastbaps), which applies the hierarchical Bayesian clustering (BHC) algorithm to determine clusters of multi-sequence genotypes (Tonkin-Hill et al., 2019). Based on the obtained alignment, 32 genes were selected based on high % of sequence conservation among available strains.

### Detection of recombinant regions

The whole genome multiple sequence alignment was analyzed in RDP5 software v. 5.5 (Martin et al., 2021).This software applies several analysis methods to identify the presence of recombinant sequences. Data were analyzed using the following recombination methods: RDP, GENECONV, Bootscan/Recscan, MaxChi, Chimaera, SiScan, and 3-seq. Recombination events considered were detected by at least 5 of these 7 methods. A recombination breakpoint graph was obtained and used to detect regions of high breakpoint concentration.

### Evolution rate and most recent common ancestor estimations

To verify the temporal signal and molecular clock likeness, the TempEst software was implemented (available at https://beast.community/tempest) providing dates of each sample. Evolutionary rate and time to the most recent common ancestor (TMRCA) were estimated through Bayesian coalescent analysis using BEAST program 10.4 version (available at https://beast.community). For this study a non-parametric Bayesian Skyride coalescent model with an uncorrelated lognormal relaxed clock was used. This analysis was run twice for 50 million Markov chain Monte Carlo (MCMC) iterations, Tracer 1.7 v. was used to check convergence evaluating that the effective sample size was higher than 200 for relevant parameters.

## Results

### Genome Characterization

The assembled genomes of ARA, CHB, HRE and NAV were 137891, 137160, 137340 and 137214bp long respectively. As expected, all four genomes contain a large central coding region surrounded by two inverted terminal repeat (ITR) regions. In each case, the left end nucleotide was designated as base 1. Aligned with other whole genome sequences of ORFV strains (Table 1), the ARA strain showed the highest similarity with SY17 strain, showing 98.7% nucleotide identity. For CHB and HRE the largest nucleotide identity was observed with NZ2, with 99.1 % and 99.0 % respectively. NAV strain showed an identity of 99.1 % with the TVL strain.

The ITRs of these viruses were 2943, 3108, 3161 and 3177 pb (ARA, CHB, HRE and NAV). The ITR of ARA contained the terminal *BamHI* sites and the telomere resolution sequence at both ends, similar to SJ1, GO, YX, MP, NA17, SY17 and CL18 strains. However, CHB, HRE and NAV strains only contained one *BamHI* site and the conserved telomere-related sequence at the right end, as the NA1 and NP strains.

The G+C nucleotide composition of these ORFV genomes were ARA 63.89%, CHB 64.22%, HRE 64.31%, NAV 64.32% which are similar to the average value of Parapoxvirus genomes. We used a moving average analysis with a 1000 bp window to analyze the G + C percentage from the genomes of the four ORFV isolates using NZ2 as a comparison (Fig. 1). The G + C percentage of each strain was predominantly high in the central region and lower in the terminal regions of the genome. Around positions 103-108 kb, a pronounced deviation from the average G + C content was observed. This recognizable signature can be seen in the other strains used in this work (Supplementary Figure 1)

**Figure 1.**
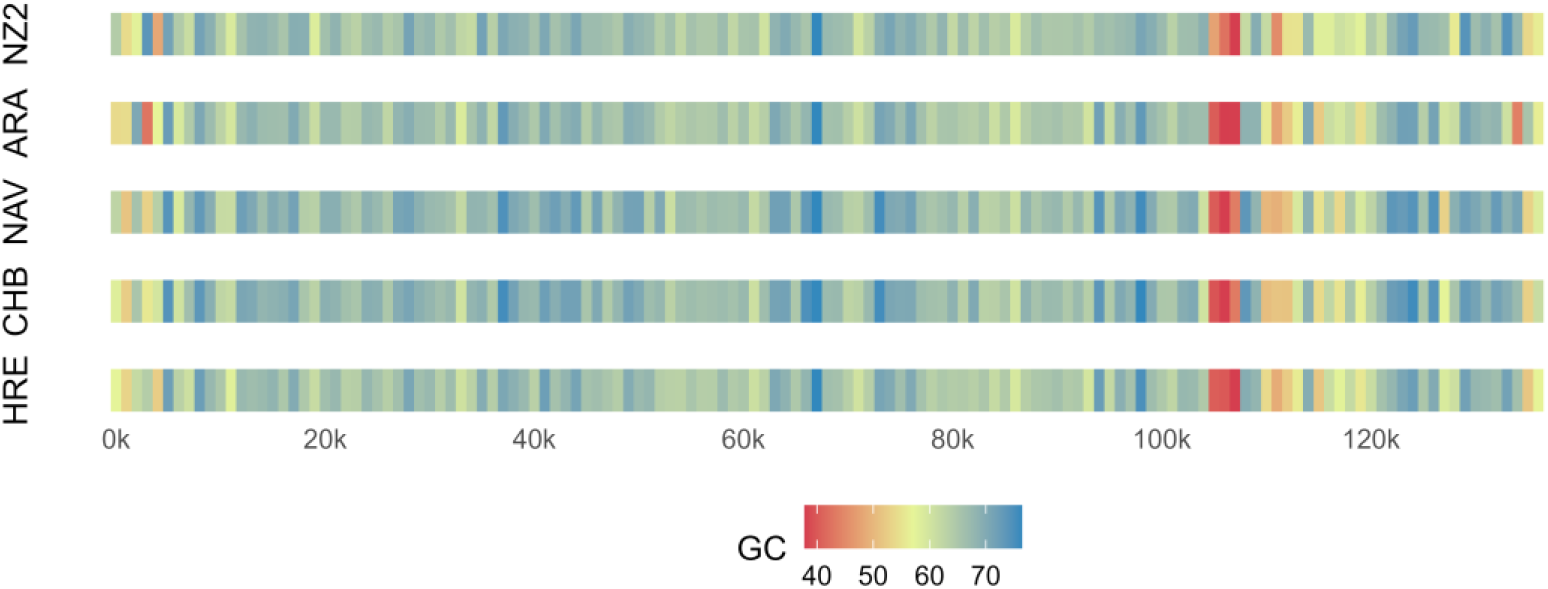
Genome Characterization of the full-length genomes. G/C content genome profile of Argentinian (HRE and CHB) and Spanish (ARA and NAV) together with reference strain ORFV-NZ2 (Sliding window size: 100 bp; made with in house R script).

Gene annotation was performed using NZ2 as reference genome, and 132 genes were identified for each sample in this study (Figure 2). Additionally, genes were predicted based on their localization within the genome, the size of the predicted proteins and by similarity with proteins previously described in Parapoxviruses. The majority of the open reading frames were non-overlapping, consistent with other poxvirus genomes and showed both, a relatively high amino acid identity and similar length with their NZ2 counterparts (Table 3).

**Figure 2.**
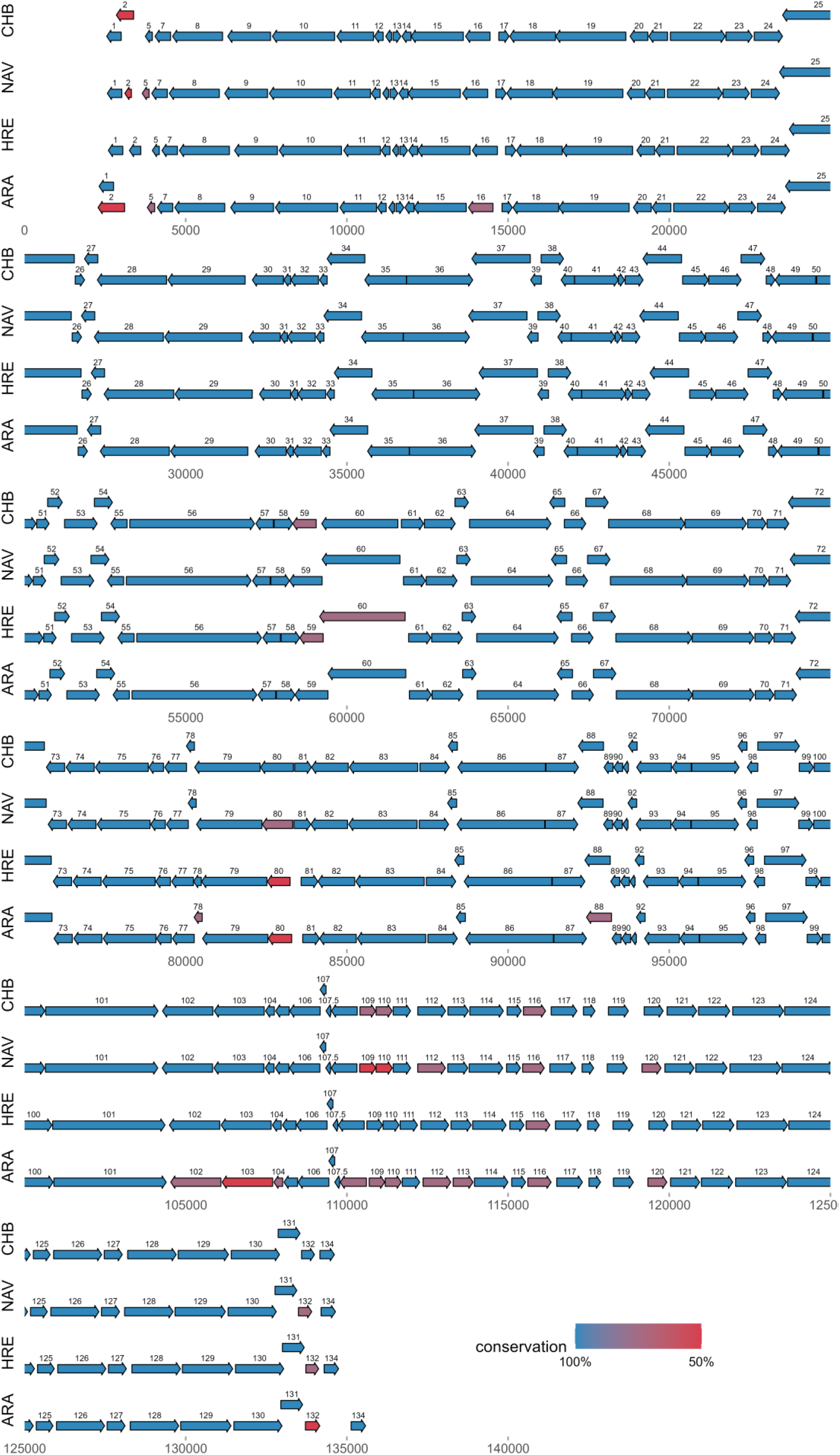
Comparative map of the CHB, HRE, ARA and NAV whole genomes. Each *orf* is represented by an arrow that indicates the size and direction of transcription; open arrowheads indicate that an *orf* is split over two lines of the diagram. Different colors of arrows indicate amino acid identity of predicted proteins between the corresponding genome and ORFV-NZ2.

Interestingly, few notable exceptions were found on the genomes described in this work, regardless the geographical origin:

1. ORF002 from ARA, which overlaps with ORF001 and is approximately 50 % larger than the encoded in NZ2.
2. ORF080 from HRE and ARA sequences, that codifies a virion core protein, showed a low identity level with respect to NZ2.
3. Similarly, the ORFs 102, 103 and 104 from ARA showed a lower level of identity with NZ2 (80, 65 and 88, respectively) than the other three isolates.
4. Meanwhile, the ORFs 109 and 110 showed lower identity in NAV isolate (64.1 and 64.4%) and somewhat higher (from 85.8 %) for ARA and CHB isolates.

### SSR and cSSR analysis

The SSR analysis of the four isolates show a Relative Abundance (RA) and a Relative Density (RD) similar to that of NZ2 (Table 2). In each case the most abundant type of repeat was the dinucleotide (DI) followed by TRI, MONO, HEXA and TETRA repeats. No PENTA repeat was detected.

**Table 2.**

| **SAMPLE** | **G+C (%)** | **SSR** | **RA** | **RD** | **MONO** | **DI** | **TRI** | **TETRA** | **PENTA** | **HEXA** | **SSR coding** | **SSR non-coding** | **cSSR** | **cSSR-coding** |
| --- | --- | --- | --- | --- | --- | --- | --- | --- | --- | --- | --- | --- | --- | --- |
| **ARA** | 83.25 | 934 | 6.77 | 48.22 | 5.35 | 75.16 | 19.06 | 0.21 | 0 | 0.21 | 90.69 | 9.31 | 64 | 93.75 |
| **NAV** | 84.00 | 946 | 6.89 | 49.32 | 5.39 | 74.84 | 19.24 | 0.21 | 0 | 0.32 | 90.80 | 9.20 | 67 | 89.55 |
| **CHB** | 83.01 | 951 | 6.93 | 49.35 | 5.47 | 75.50 | 18.61 | 0.00 | 0 | 0.42 | 90.64 | 9.36 | 64 | 87.50 |
| **HRE** | 84.15 | 936 | 6.82 | 48.78 | 5.34 | 74.68 | 19.34 | 0.32 | 0 | 0.32 | 91.77 | 8.23 | 63 | 93.65 |
| **NZ2** | 83.54 | 954 | 6.92 | 49.64 | 5.24 | 74.84 | 19.08 | 0.31 | 0 | 0.52 | 90.67 | 9.33 | 68 | 86.76 |
| **SA00** | 80.64 | 1012 | 7.23 | 51.96 | 4.94 | 74.51 | 18.28 | 2.08 | 0 | 0.20 | 85.08 | 14.92 | 75 | 82.67 |

SSR. The counts of microsatellites in each genome.
RA. Relative abundance of microsatellites (number of microsatellites or compound microsatellites/kb).
RD. Relative density of microsatellites is defined as the total length (bp) contributed by each microsatellite per kb of sequence.
MONO, DI, TETRA, PENTA and HEXA are the percentage of each type of repeat in each genome.
SSR coding and SSR non-coding are the percentage of SSR in genic and intergenic regions, respectively, for each genome.
cSSR. The counts of compound microsatellites in each genome.

**Table 3.**
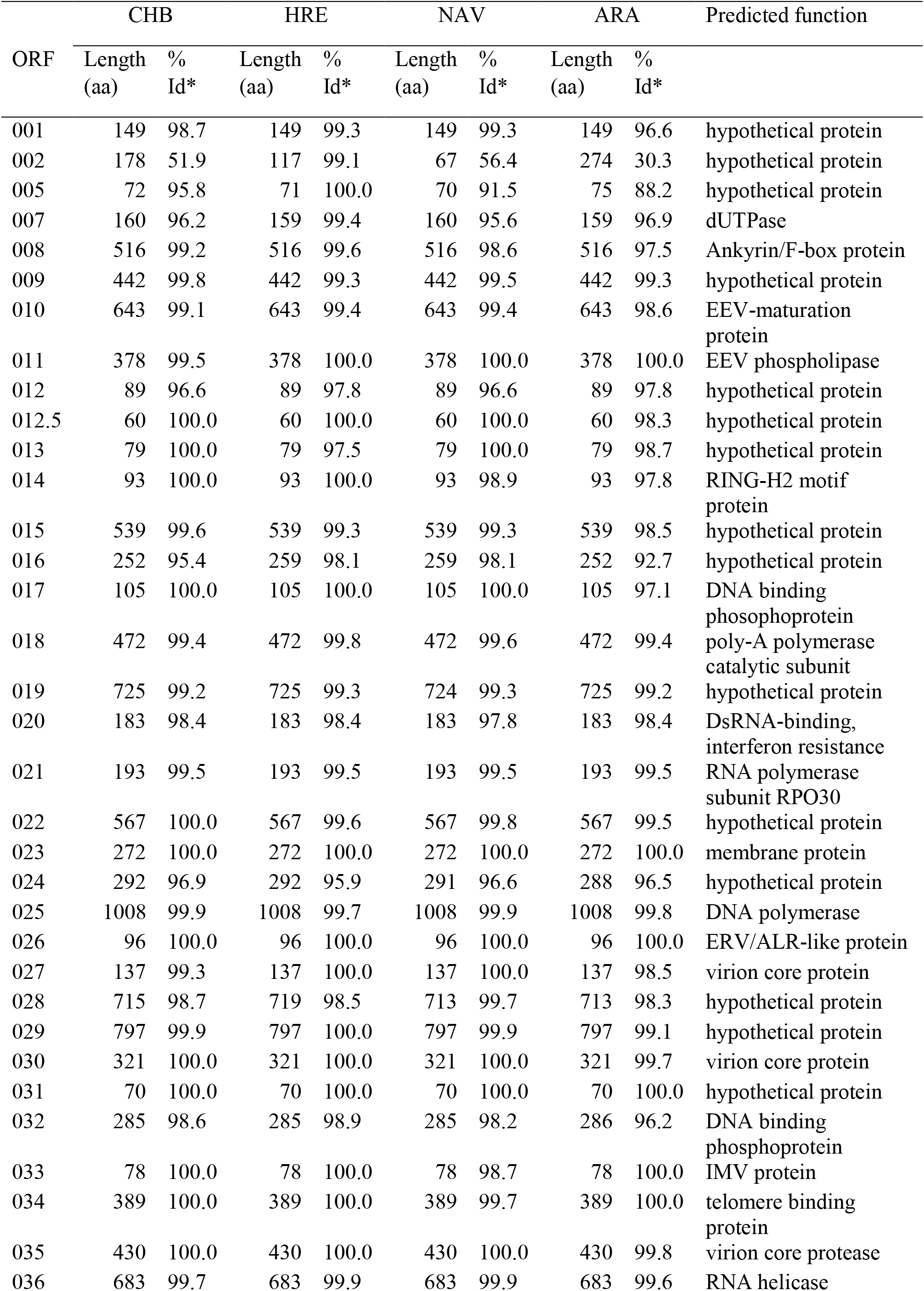
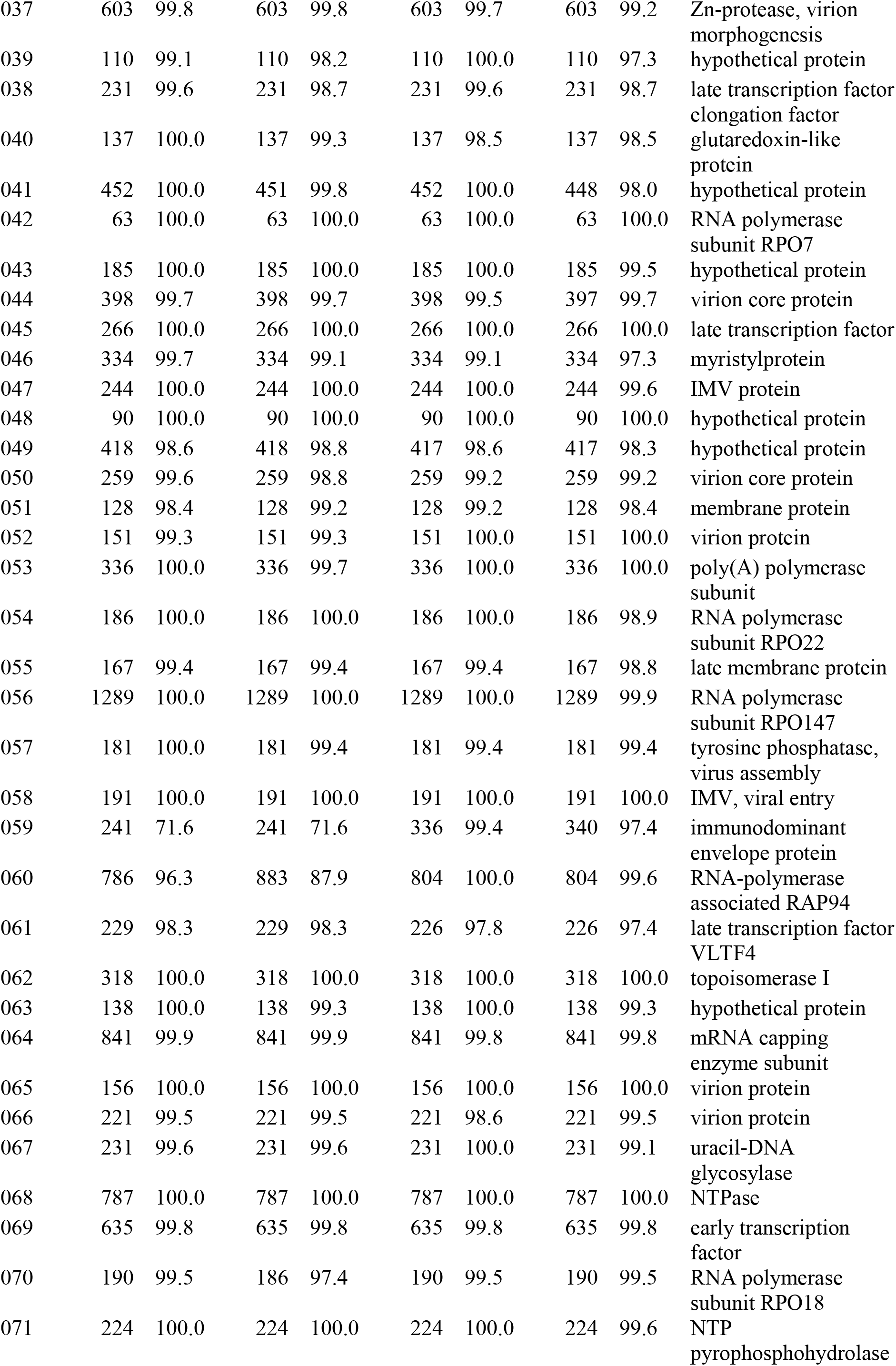
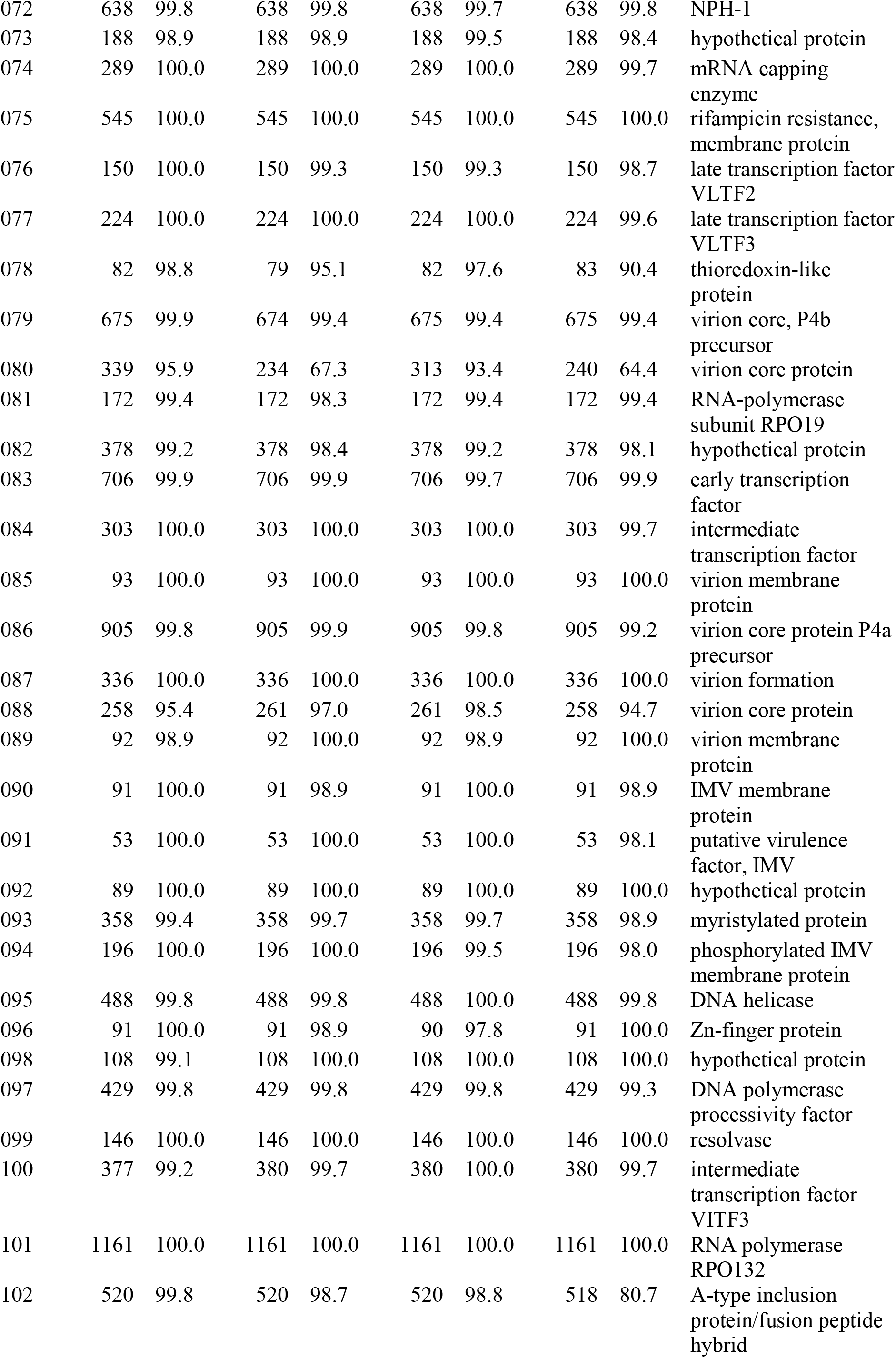
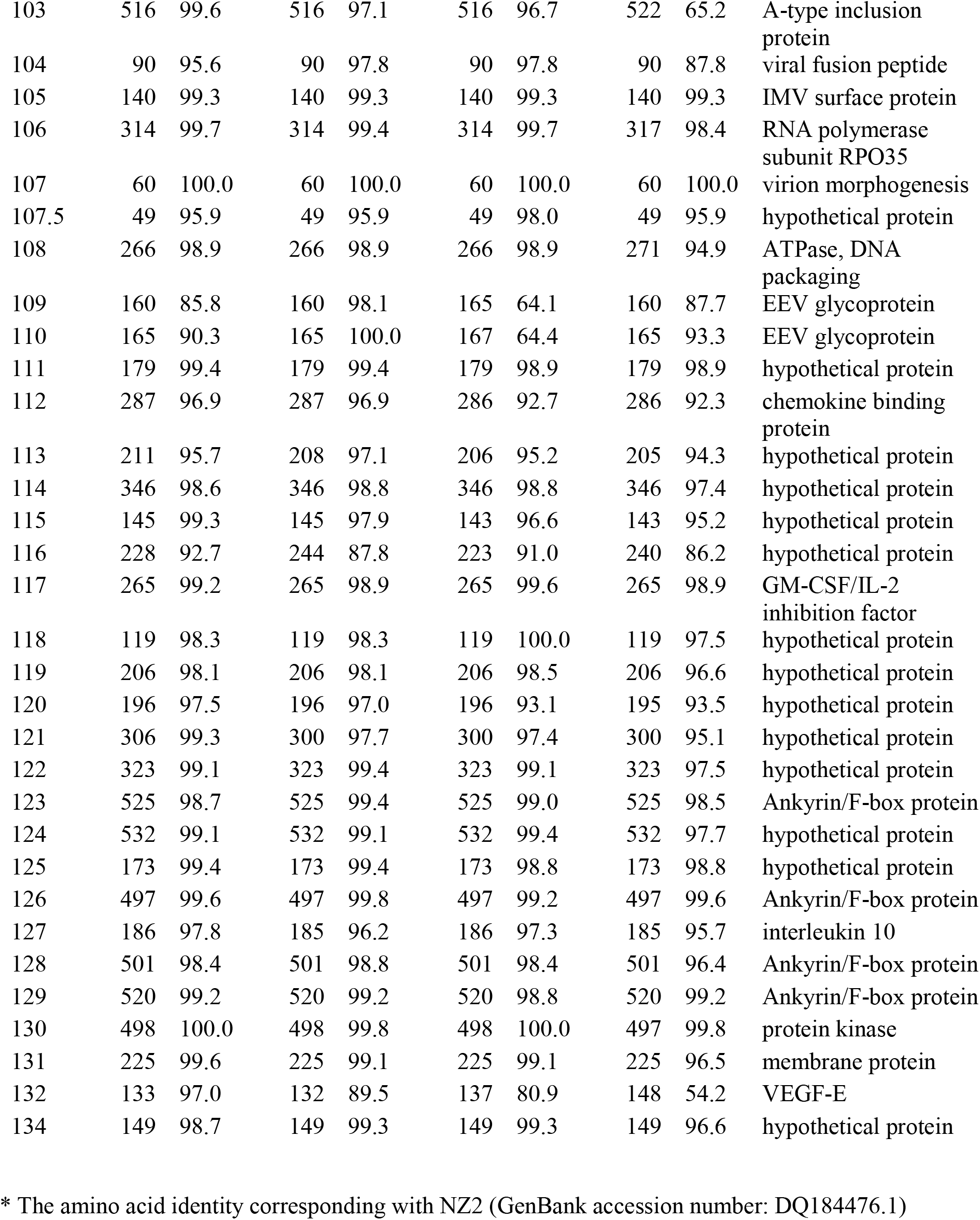
ORFs of CHB, HRE, NAV y ARA compared with the corresponding ORFs of NZ2 strain.

**Table 4.**
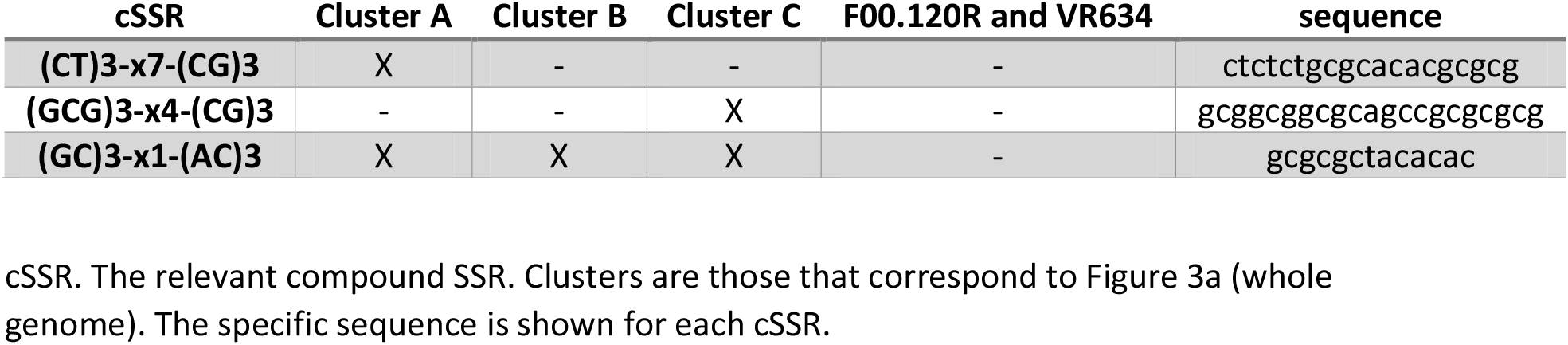

Additionally, the compound SSR (cSSR) composition was obtained and the occurrence of each cSSR in the 21 Parapoxvirus full genomes used in this work was analyzed. Of the 308 cSSR found, more than one half (159) were present in only one sample each. Furthermore, 19 ORFV complete sequences presented the cSSR (GC)3-x1-(AC)3, that was not present in PCPV samples (F00.120R and VR634). Interestingly, cluster A and cluster C presented a specific cSSR each, (CT)3-x7-(CG)3 and (GCG)3-x4-(CG)3, respectively.

### Phylogenetic analysis

The phylogenetic analysis was performed at three different levels: full genome, excluding breakpoint regions and using 32 selected genes (Figure 3 and supplementary figure 2).

**Figure 3.**
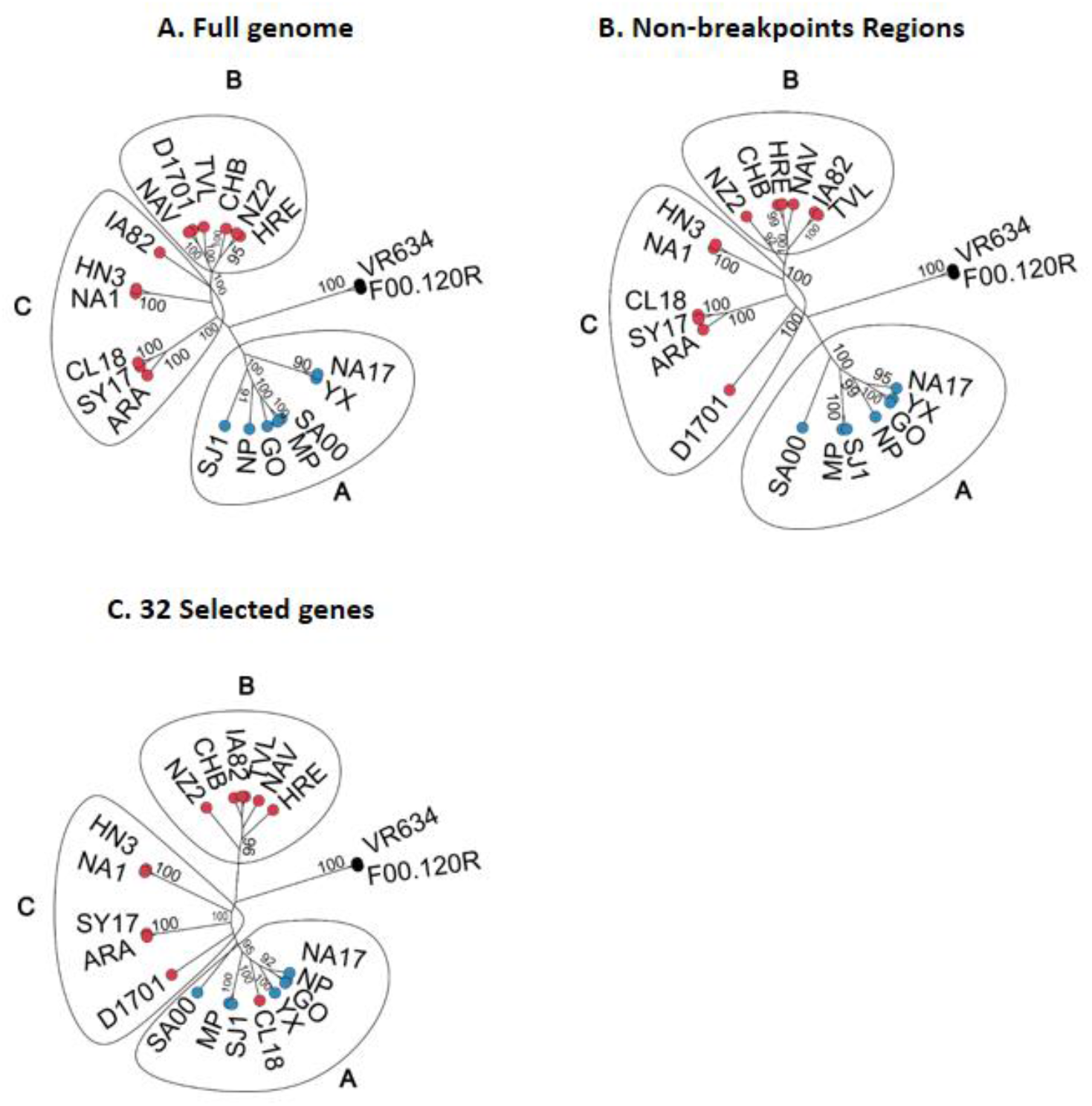
Phylogenetic analysis. Maximum-Likelihood phylogenetic tree based on full-length genome sequences(A); genome sequences, excluding breakpoints (B); and 32 concatenated selected genes (C). The numbers at nodes correspond to bootstrap values. Tip nodes are colored in blue or red for goat and sheep samples, respectively.

The phylogenetic tree derived from the full-length genome alignment was well supported, with a group structure of three clusters, namely A, B and C (Figure 3a).

Cluster A is composed mostly by sequences from Chinese origin, with the exceptions of SA00 sample that comes from United States, and MP sample from India.

Meanwhile, cluster B grouped sequences of different countries like Argentina, Spain, New Zealand, United States and Germany. Finally, cluster C was composed by Chinese samples, one Spanish sample and one USA sample (Figure 3a). Three of the samples sequenced and analyzed in this work (CHB, NAV and HRE) showed a close phylogenetic relationship within cluster B, while the ARA sample was more related to the Chinese samples in cluster C.

Interestingly, we observed a clear separation between species, with goat samples exclusively on cluster A and the sheep samples distributed across two clusters (B and C). In order to test if a similar separation between species could be obtained by using less than whole genome data, high breakpoint concentration regions, were detected and excluded from the alignment (positions 19000 to 100904 and 116402 to 151219 of the full-length genome alignment were maintained). A 116721 bp-long concatenated sequence was obtained for each sample. The obtained tree showed a similar group structure with the three clusters (Figure 3b). However, sample D1701 shifted from clade B to clade C and sample IA82 from clade C to clade B.

Moreover, we selected 32 highly conserved genes for the family Poxviridae (Babkin and Babkina, 2011), all present in the dataset excluding breakpoint regions. Specifically, a 42417 bp-long concatenated sequence was identified and the obtained tree was well supported (bootstrap value of 100%). The maximum likelihood tree showed a similar group structure, with the tree clusters mostly composed of individuals of the same species, goat (A) or sheep (B and C) with the exception of the sheep sample CL18 present in the goat cluster A (Figure 3c). Of note, root location in phylograms shifted towards the middle of cluster C.

### Evolutionary estimated rate and most recent common ancestor

We performed a molecular clock analysis on all three datasets obtaining well-supported Bayesian trees (posterior probability of 1), confirming clade composition and group structure observed with maximum-likelihood method (data not shown). No convergence was obtained when full-length genome or the non-breakpoint regions datasets were considered. But convergence was obtained for the 32 selected genes dataset.

The temporal signal analysis exhibited a positive correlation between genetic divergence and time, however diffuse regression plots (R^2^=3.16e-3) were obtained using the full-length genome dataset (Supplementary File 1). Therefore, two bigger groups (A and B+C) were analyzed independently. Using this approach, we also found a positive correlation between the two variables, and a slight improvement in the regression coefficient for cluster A (R^2^=6.61e-2) and B/C (R^2^=0.1). These results strongly suggest an uneven evolutionary rate among the branches, so a relaxed molecular clock was chosen for the next phylogenetic analysis.

The MCMC analysis, performed with Beast software, showed that the nucleotide substitution rate for this dataset was 3.2×10^-5^ subs/site/year (95% HPD 4.2X10^-8^ -7.6 X10^-5^) with a median value of 2.8 × 10^-5^ subs/site/year. The TMRCA placed the median divergence time in 1779 (HPD interval 95%: 499, 1967).

## DISCUSSION

In this work, we isolated and sequenced four novel Orf virus strains named ARA, CHB, HRE and NAV from infected sheep in Argentina and Spain. The whole genomic sequence of the isolates ranged between 137160 and 137891 bp, including the inverted terminal repeats. Despite differences in length, all the sequences obtained resembled those publicly available in gene composition, encoding 132 non-overlapping genes as featured by other Orf viruses.

The G + C content for all samples was 64% approximately. This value coincides with that described for other genomes of the Parapoxvirus (PPV) genus (Mercer et al., 2006). A small region located approximately between 103-108 kb with a low G+C concentration (40-45%) draws attention. This coincides with previous findings in ORFV genomes, and also in PCPV and BPSV genomes, so it could be considered a molecular signature of the PPV genus as previously proposed (Hautaniemi et al., 2010; Mercer et al., 2006).

Characterization of the four genomes with respect to microsatellites showed between 934 and 951 SSRs per genome, with RA and RD values between 6.77-6.89 and 48.22-49.35 respectively. These values are slightly lower than those reported by Sahu et al., 2020, but within the ranges found for other DNA viruses (Ouyang et al., 2012; Singh et al., 2014). In the four genomes, the most abundant repetitive units were: dinucleotide (75.04%), trinucleotide (19.06%) and mononucleotide (5.38%). The majority of the SSRs and cSSRs were found in coding regions (90.97% and 91.11% respectively). This distribution is reasonable since viruses have short intergenic regions and can often overlap with coding regions.

Certain studies have suggested the use of SSR and cSSR for the characterization of viral strains (Burrel et al., 2013; Houng et al., 2009). We focused on the analysis of cSSRs since, given the complexity of their structure, they could serve as specific spots to determine relationships between viral isolates. Cluster A (goat) and cluster C (sheep) (Figure 3) presented a specific cSSR each: (CT)3-x7-(CG)3 and (GCG)3-x4-(CG)3, respectively. It was not possible to find a cSSR as a marker for group B. Maintaining the structure of cSSRs is complex because different forces act: mutations, duplication, recombination, etc. It is possible that the members of group B have an incomplete cSSR and therefore cannot be detected by standard microsatellite studies (Delgrange and Rivals, 2004; Mudunuri and Nagarajaram, 2007). Having information from more genomes would allow us to determine if these cSSRs are really a molecular signature of groups A and C or if they are a residual in the differentiation of these groups.

Our phylogenetic analyses with full-length genomes (Figure 3A), and in datasets with partial sequences, based on the whole genome excluding high breakpoint concentration regions (Figure 3B) and the 32 selected conserved genes (Figure 3C) suggest a viral population structuring in three well-supported clusters.

The four samples that originated from sheep and were isolated from Argentina (CHB and HRE) and Spain (NAV and ARA) grouped with other sheep samples. A phylogenetic analysis based on the whole genomic sequences of 19 ORFV strains revealed that CHB and HRE strains isolated from Argentinian sheep have a close relationship with NZ2 from New Zealand, while Spanish sequence NAV was highly similar to North American TVL strain, and ARA (Spain) was associated to Chinese SY17 and CL18 strains.

Regarding the two Spanish samples sequenced and analyzed in this paper, it is remarkable that both belong to two different groups/clades, confirming the existence of at least two genomic clusters circulating in this country. On the contrary, the two Argentinean samples showed a closer relationship within the same cluster.

Nevertheless, previous descriptions have found the presence of distinct clusters in the country (Peralta et al., 2018) based on analysis of partial sequences from genes orf011, orf020, orf109, and orf127, so it is essential to gather data from additional sequences in order to accept or reject this hypothesis. In addition to the difference in the data set that was used in both studies, in Peralta et al 2018, ORFV strains from goats were analyzed, so we could be observing a different evolutionary history of the Orf virus in Argentina, depending on the host species. However, it is essential to gather data from additional sequences in order to accept or reject this hypothesis.

Interestingly, some research using specific genes (Chi et al., 2015; Coradduzza et al., 2021) has established a clustering of sequences by host. Genomic culstering according to host species could be found in our analysis. However, assessment of sequences to groups A and B+C could be due to geographic location, the host species or both. Additional analysis including more sequences would clarify this point.

Due to the size and complexity of the genome, the analysis using whole genome sequences can generate discordant signals and incompatible results when compared to partial sequence analysis. For this reason, in this work, we conducted a gene selection reducing the interfering signals, bioinformatics cost and offering more reproducible results. The 32 selected genes were selected, not only because they are highly conserved among Poxviruses, but also because their orthologs in Orthopoxviruses were shown to have the same evolutionary rate (Babkin and Babkina, 2011). It is not surprising that only with this set of data, it was possible to reach convergence and obtain molecular dating of the analyzed sequences.

As far as we know, this is the first molecular epidemiology study that includes ORFV sequences using 32 highly conserved genes and showing comparable results to the full-length genome.

The nucleotide substitution rate inferred in this work was 3.2×10^-5^ subs/site/year. This value could be considered high for a DNA virus, but in accordance with previous works studying Myxoma virus (Kerr et al., 2017) or Avipox viruses (Le Loc’h et al., 2015). Noteworthy, the analysis of MCRA suggested a date in the late XVII century, not far from the first case of ORFV reported in a sheep by Steeb in 1787 (Barraviera, 2005). However, wide error margins were observed, likely due to the low number of sequences. Greater precision could be expected in dating the common ancestor by incorporating members of the Parapoxvirus genus. However, very few PCPV and BPSV genomes are available.

Since sheep and goats are not native animals to South America, it could be hypothesized that the Orf virus arrived on the continent with colonization from Europe. The analysis of the four genomes available in our study (two from Spain and two from Argentina) does not allow us to confirm this hypothesis, although the Spanish NAV strain belongs to the same group as the Argentine HRE and CHB. While whole-genome analysis can be challenging, it provides much more information than analysis of individual genes, where each gene can tell a different evolutionary story (Li et al., 2023).

In this work, we determined the complete genome of two ORFV strains from Spain and two from Argentina, the first report being for South America and the second for Europe. Genetic characterization of ORFV strains is the first step towards the development of molecular tools oriented to diagnostics and vaccine development. Furthermore, increasing the knowledge on ORFV strains genetic composition will establish relationships and contribute to future epidemiological studies. Moreover, the ORFV evolutionary analysis will gain precision when more whole genome sequences become available.

## Acknowledgements

Samples from diseased animals were obtained and kindly provided by INTIA from Navarra and by Hector Ruiz (DVM) from Aragon.

Argentine samples were obtained by Carlos Rossanigo (INTA-San Luis) and Alejandro Varela (Universidad Nacional de La Plata), HRE and CHB respectively. Part of the results were obtained thanks to the following grants: PICT2017-2728 from Agencia de Promoción Científica y Tecnológica, Argentina.

## Notes

### Competing Interest Statement

The authors have declared no competing interest.

## Bibliography

Alam, C.M., Singh, A.K., Sharfuddin, C., Ali, S., 2013. In-silico analysis of simple and imperfect microsatellites in diverse tobamovirus genomes. Gene 530, 193–200. 10.1016/j.gene.2013.08.046

Babkin, I.V., Babkina, I.N., 2011. Molecular dating in the evolution of vertebrate poxviruses. Intervirology 54, 253–260.

Bankevich, A., Nurk, S., Antipov, D., Gurevich, A.A., Dvorkin, M., Kulikov, A.S., Lesin, V.M., Nikolenko, S.I., Pham, S., Prjibelski, A.D., Pyshkin, A.V., Sirotkin, A.V., Vyahhi, N., Tesler, G., Alekseyev, M.A., Pevzner, P.A., 2012. SPAdes: A New Genome Assembly Algorithm and Its Applications to Single-Cell Sequencing. J. Comput. Biol. 19, 455–477. 10.1089/cmb.2012.0021

Barraviera, S., 2005. Diseases caused by poxvirus-orf and milker’s nodules: a review. J. Venom. Anim. Toxins Trop. Dis. 11, 102–108.

Burrel, S., Ait-Arkoub, Z., Voujon, D., Deback, C., Abrao, E.P., Agut, H., Boutolleau, D., 2013. Molecular Characterization of Herpes Simplex Virus 2 Strains by Analysis of Microsatellite Polymorphism. J. Clin. Microbiol. 51, 3616–3623. 10.1128/JCM.01714-13

Burriel, A.R., 1997. Udder Orf Infection and its Role in Ovine Clinical Mastitis Caused by Pasteurella haemolytica. J. Trace Elem. Med. Biol. 11, 28–31. 10.1016/S0946-672X(97)80006-5

Bushnell, 2019. BBMap [WWW Document]. SourceForge. URL https://sourceforge.net/projects/bbmap/ (accessed 6.26.24).

Chan, K.-W., Lin, J.-W., Lee, S.-H., Liao, C.-J., Tsai, M.-C., Hsu, W.-L., Wong, M.-L., Shih, H.-C., 2007. Identification and phylogenetic analysis of orf virus from goats in Taiwan. Virus Genes 35, 705–712. 10.1007/s11262-007-0144-6

Chen, M., Tan, Z., Jiang, J., Li, M., Chen, H., Shen, G., Yu, R., 2009. Similar distribution of simple sequence repeats in diverse completed *Human Immunodeficiency Virus Type 1* genomes. FEBS Lett. 583, 2959–2963. 10.1016/j.febslet.2009.08.004

Chi, X., Zeng, X., Li, W., Hao, W., Li, M., Huang, X., Huang, Y., Rock, D.L., Luo, S., Wang, S., 2015. Genome analysis of orf virus isolates from goats in the Fujian Province of southern China. Front. Microbiol. 6. 10.3389/fmicb.2015.01135

Coradduzza, E., Sanna, D., Rocchigiani, A.M., Pintus, D., Scarpa, F., Scivoli, R., Bechere, R., Dettori, M.A., Montesu, M.A., Marras, V., Lobrano, R., Ligios, C., Puggioni, G., 2021. Molecular Insights into the Genetic Variability of ORF Virus in a Mediterranean Region (Sardinia, Italy). Life 11. 10.3390/life11050416

Danecek, P., Bonfield, J.K., Liddle, J., Marshall, J., Ohan, V., Pollard, M.O., Whitwham, A., Keane, T., McCarthy, S.A., Davies, R.M., Li, H., 2021. Twelve years of SAMtools and BCFtools. GigaScience 10, giab008. 10.1093/gigascience/giab008

Davis, C.L., Field, D., Metzgar, D., Saiz, R., Morin, P.A., Smith, I.L., Spector, S.A., Wills, C., 1999. Numerous Length Polymorphisms at Short Tandem Repeats in Human Cytomegalovirus. J. Virol. 73, 6265–6270. 10.1128/JVI.73.8.6265-6270.1999

Delgrange, O., Rivals, E., 2004. STAR: an algorithm to Search for Tandem Approximate Repeats. Bioinformatics 20, 2812–2820. 10.1093/bioinformatics/bth335

Delhon, G., Tulman, E.R., Afonso, C.L., Lu, Z., De La Concha-Bermejillo, A., Lehmkuhl, H.D., Piccone, M.E., Kutish, G.F., Rock, D.L., 2004. Genomes of the Parapoxviruses Orf Virus and Bovine Papular Stomatitis Virus. J. Virol. 78, 168–177. 10.1128/JVI.78.1.168-177.2004

Guo, J., Rasmussen, J., Wünschmann, A., De La Concha-Bermejillo, A., 2004. Genetic characterization of orf viruses isolated from various ruminant species of a zoo. Vet. Microbiol. 99, 81–92. 10.1016/j.vetmic.2003.11.010

Haig, D.M., Mercer, A.A., 1998. Ovine diseases. Orf. Vet. Res. 29, 311–326.

Hatcher, E., Wang, C., Lefkowitz, E., 2015. Genome Variability and Gene Content in Chordopoxviruses: Dependence on Microsatellites. Viruses 7, 2126–2146. 10.3390/v7042126

Hautaniemi, M., Ueda, N., Tuimala, J., Mercer, A.A., Lahdenpera, J., McInnes, C.J., 2010. The genome of pseudocowpoxvirus: comparison of a reindeer isolate and a reference strain. J. Gen. Virol. 91, 1560–1576. 10.1099/vir.0.018374-0

Hosamani, M., Bhanuprakash, V., Scagliarini, A., Singh, R.K., 2006. Comparative sequence analysis of major envelope protein gene (B2L) of Indian orf viruses isolated from sheep and goats. Vet. Microbiol. 116, 317–324. 10.1016/j.vetmic.2006.04.028

Houng, H.-S.H., Lott, L., Gong, H., Kuschner, R.A., Lynch, J.A., Metzgar, D., 2009. Adenovirus Microsatellite Reveals Dynamics of Transmission during a Recent Epidemic of Human Adenovirus Serotype 14 Infection. J. Clin. Microbiol. 47, 2243– 2248. 10.1128/JCM.01659-08

Karki, M., Kumar, A., Arya, S., Ramakrishnan, M.A., Venkatesan, G., 2019. Poxviral E3L ortholog (Viral Interferon resistance gene) of orf viruses of sheep and goats indicates species-specific clustering with heterogeneity among parapoxviruses. Cytokine 120, 15–21. 10.1016/j.cyto.2019.04.001

Katoh, K., Standley, D.M., 2013. MAFFT Multiple Sequence Alignment Software Version 7: Improvements in Performance and Usability. Mol. Biol. Evol. 30, 772–780. 10.1093/molbev/mst010

Kerr, P.J., Cattadori, I.M., Rogers, M.B., Fitch, A., Geber, A., Liu, J., Sim, D.G., Boag, B., Eden, J.-S., Ghedin, E., 2017. Genomic and phenotypic characterization of myxoma virus from Great Britain reveals multiple evolutionary pathways distinct from those in Australia. PLoS Pathog. 13, e1006252.

Kottaridi, C., Nomikou, K., Lelli, R., Markoulatos, P., Mangana, O., 2006. Laboratory diagnosis of contagious ecthyma: Comparison of different PCR protocols with virus isolation in cell culture. J. Virol. Methods 134, 119–124. 10.1016/j.jviromet.2005.12.005

Le Loc’h, G., Bertagnoli, S., Ducatez, M.F., 2015. Time scale evolution of avipoxviruses. Infect. Genet. Evol. 35, 75–81. 10.1016/j.meegid.2015.07.031

Li, S., Jing, T., Zhu, F., Chen, Y., Yao, X., Tang, X., Zuo, C., Liu, M., Xie, Y., Jiang, Y., Wang, Y., Li, D., Li, L., Gao, S., Chen, D., Zhao, H., Ma, W., 2023. Genetic Analysis of Orf Virus (ORFV) Strains Isolated from Goats in China: Insights into Epidemiological Characteristics and Evolutionary Patterns. Virus Res. 334, 199160. 10.1016/j.virusres.2023.199160

Martin, D.P., Varsani, A., Roumagnac, P., Botha, G., Maslamoney, S., Schwab, T., Kelz, Z., Kumar, V., Murrell, B., 2021. RDP5: a computer program for analyzing recombination in, and removing signals of recombination from, nucleotide sequence datasets. Virus Evol. 7, veaa087. 10.1093/ve/veaa087

Mercer, A.A., Ueda, N., Friederichs, S.-M., Hofmann, K., Fraser, K.M., Bateman, T., Fleming, S.B., 2006. Comparative analysis of genome sequences of three isolates of Orf virus reveals unexpected sequence variation. Virus Res. 116, 146–158. 10.1016/j.virusres.2005.09.011

Minh, B.Q., Schmidt, H.A., Chernomor, O., Schrempf, D., Woodhams, M.D., Von Haeseler, A., Lanfear, R., 2020. IQ-TREE 2: New Models and Efficient Methods for Phylogenetic Inference in the Genomic Era. Mol. Biol. Evol. 37, 1530–1534. 10.1093/molbev/msaa015

Mrázek, J., Guo, X., Shah, A., 2007. Simple sequence repeats in prokaryotic genomes. Proc. Natl. Acad. Sci. 104, 8472–8477. 10.1073/pnas.0702412104

Mudunuri, S.B., Kumar, P., Rao, A.A., Pallamsetty, S., Nagarajaram, H.A., 2010. G-IMEx: A comprehensive software tool for detection of microsatellites from genome sequences. Bioinformation 5, 221–223. 10.6026/97320630005221

Mudunuri, S.B., Nagarajaram, H.A., 2007. IMEx: Imperfect Microsatellite Extractor. Bioinformatics 23, 1181–1187. 10.1093/bioinformatics/btm097

Peralta, A., Robles, C.A., Micheluod, J.F., Rossanigo, C.E., Martinez, A., Carosio, A., König, G.A., 2018. Phylogenetic Analysis of ORF Viruses From Five Contagious Ecthyma Outbreaks in Argentinian Goats. Front. Vet. Sci. 5, 134. 10.3389/fvets.2018.00134

Rambaut, A., Drummond, A.J., Xie, D., Baele, G., Suchard, M.A., 2018. Posterior Summarization in Bayesian Phylogenetics Using Tracer 1.7. Syst. Biol. 67, 901–904. 10.1093/sysbio/syy032

Ronquist, F., Teslenko, M., Van Der Mark, P., Ayres, D.L., Darling, A., Höhna, S., Larget, B., Liu, L., Suchard, M.A., Huelsenbeck, J.P., 2012. MrBayes 3.2: Efficient Bayesian Phylogenetic Inference and Model Choice Across a Large Model Space. Syst. Biol. 61, 539–542. 10.1093/sysbio/sys029

Sahu, B.P., Majee, P., Singh, R.R., Sahoo, A., Nayak, D., 2020. Comparative analysis, distribution, and characterization of microsatellites in Orf virus genome. Sci. Rep. 10, 13852. 10.1038/s41598-020-70634-6

Tcherepanov, V., Ehlers, A., Upton, C., 2006. Genome Annotation Transfer Utility (GATU): rapid annotation of viral genomes using a closely related reference genome. BMC Genomics 7, 150. 10.1186/1471-2164-7-150

Tonkin-Hill, G., Lees, J.A., Bentley, S.D., Frost, S.D.W., Corander, J., 2019. Fast hierarchical Bayesian analysis of population structure. Nucleic Acids Res. 47, 5539–5549. 10.1093/nar/gkz361

Tóth, G., Gáspári, Z., Jurka, J., 2000. Microsatellites in Different Eukaryotic Genomes: Survey and Analysis. Genome Res. 10, 967–981. 10.1101/gr.10.7.967

Velazquez-Salinas, L., Ramirez-Medina, E., Bracht, A.J., Hole, K., Brito, B.P., Gladue, D.P., Carrillo, C., 2018. Phylodynamics of parapoxvirus genus in Mexico (2007–2011). Infect. Genet. Evol. 65, 12–14. 10.1016/j.meegid.2018.07.005

Walker, A., Petheram, S.J., Ballard, L., Murph, J.R., Demmler, G.J., Bale, J.F., 2001. Characterization of Human Cytomegalovirus Strains by Analysis of Short Tandem Repeat Polymorphisms. J. Clin. Microbiol. 39, 2219–2226. 10.1128/JCM.39.6.2219-2226.2001

Wu, X., Zhou, L., Zhao, X., Tan, Z., 2014. The analysis of microsatellites and compound microsatellites in 56 complete genomes of Herpesvirales. Gene 551, 103–109. 10.1016/j.gene.2014.08.054

Yao, X., Pang, M., Wang, T., Chen, X., Tang, X., Chang, J., Chen, D., Ma, W., 2020. Genomic Features and Evolution of the Parapoxvirus during the Past Two Decades. Pathogens 9, 888. 10.3390/pathogens9110888

Zwartouw, H.T., Westwood, J.C.N., Appleyard, G., 1962. Purification of Pox Viruses by Density Gradient Centrifugation. J. Gen. Microbiol. 29, 523–529. 10.1099/00221287-29-3-523

