## Supplementary files for "The whole genome analysis of four Orf virus strains from Europe and South America"

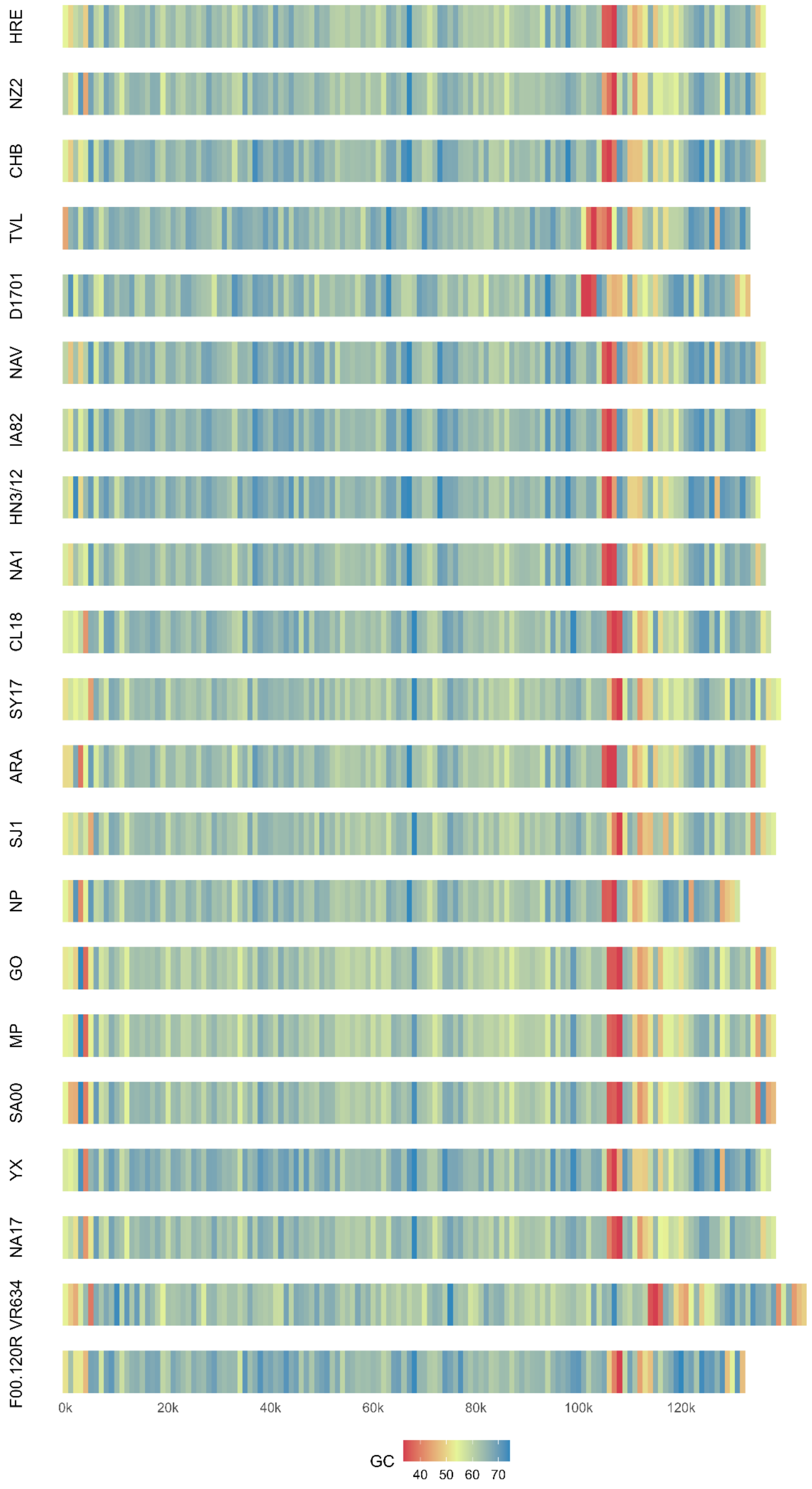
HRE
NZ2
CHB
TVL
D1701
NAV
IA82
HN3/12
NA1
CL18
SY17
ARA
SJ1
NP
GO
MP
SA00
YX
NA17
VR634
F00.120R
0k
20k
40k
60k
80k
100k
120k
GC
40
50
60
70

**Supplementary Figure 1.** G/C genome profile of all 21 genomes used in this study (Sliding window size: 100 bp; made with in house R script).

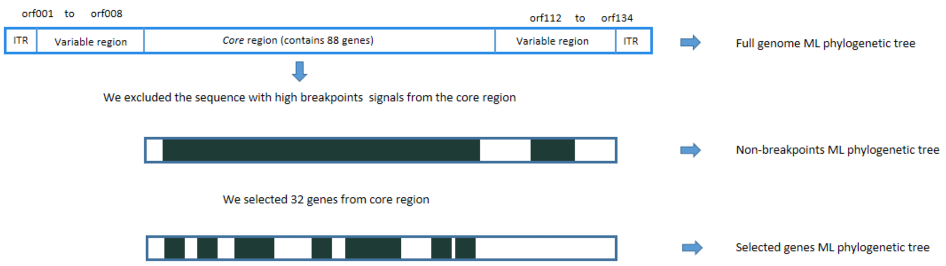


**Supplementary Figure 2.** Schematic representation of the genome regions used for the phylogenetic analysis. Diagram is not at scale. The genes selected for the analysis were: 011, 018, 021, 025, 036, 038, 040, 042, 045, 053, 054, 056, 057, 060, 061, 062, 064, 067, 068, 069, 070, 072, 074, 076, 077, 079, 081, 083, 084, 086, 095, 096.

### Supplementary File 1

#### TEMPEST

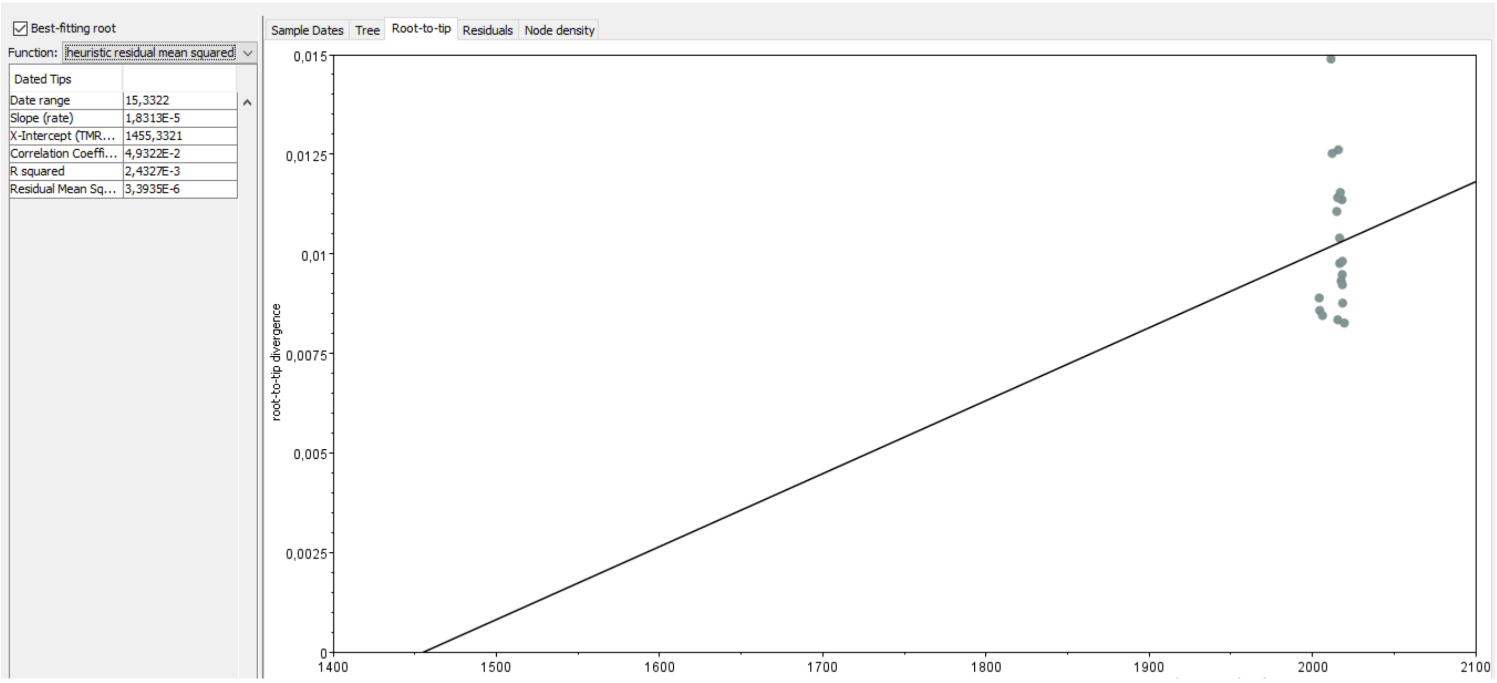


#### Combined TRACER

#### AGE_ROOT

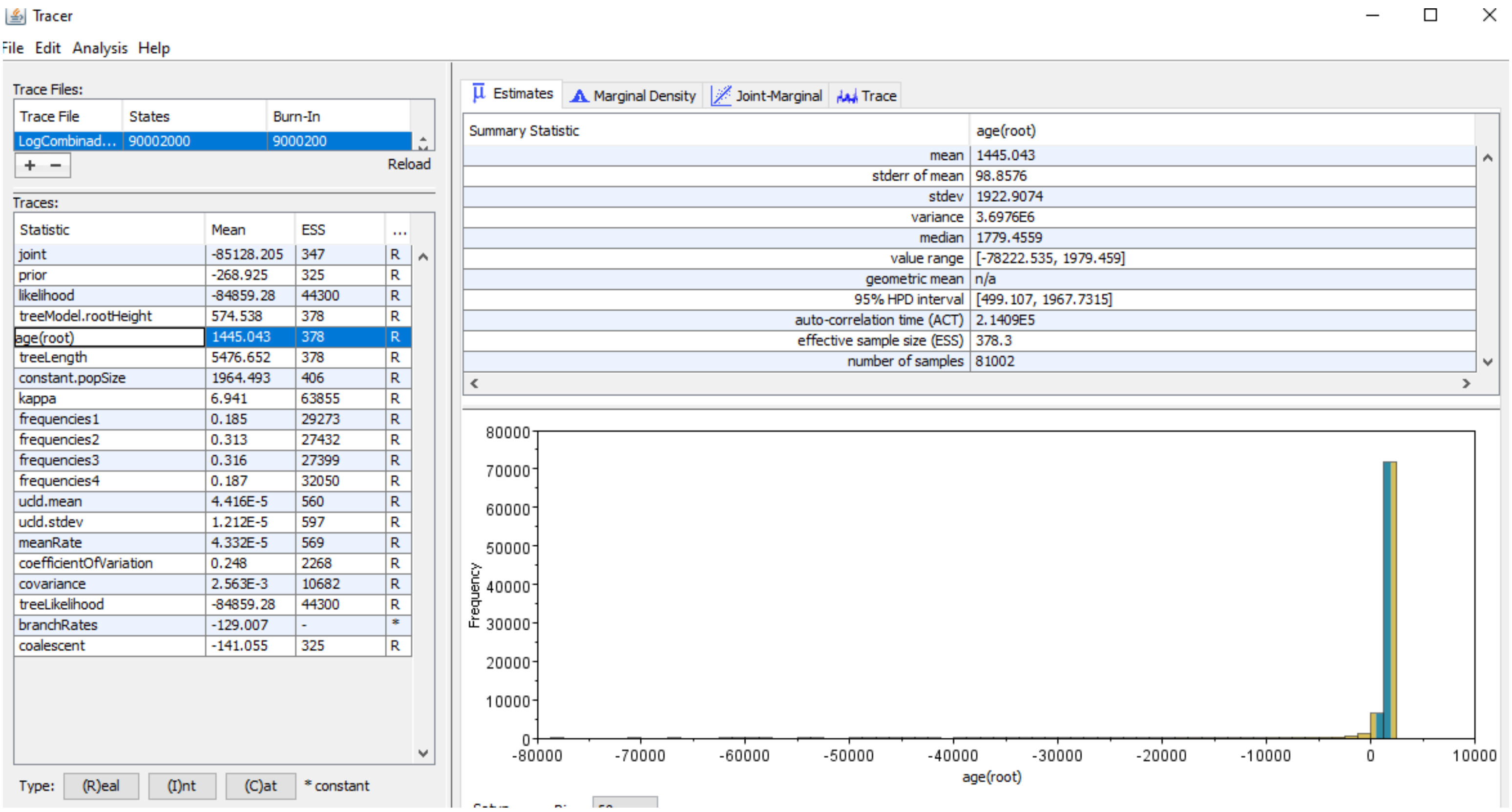


**RATE**

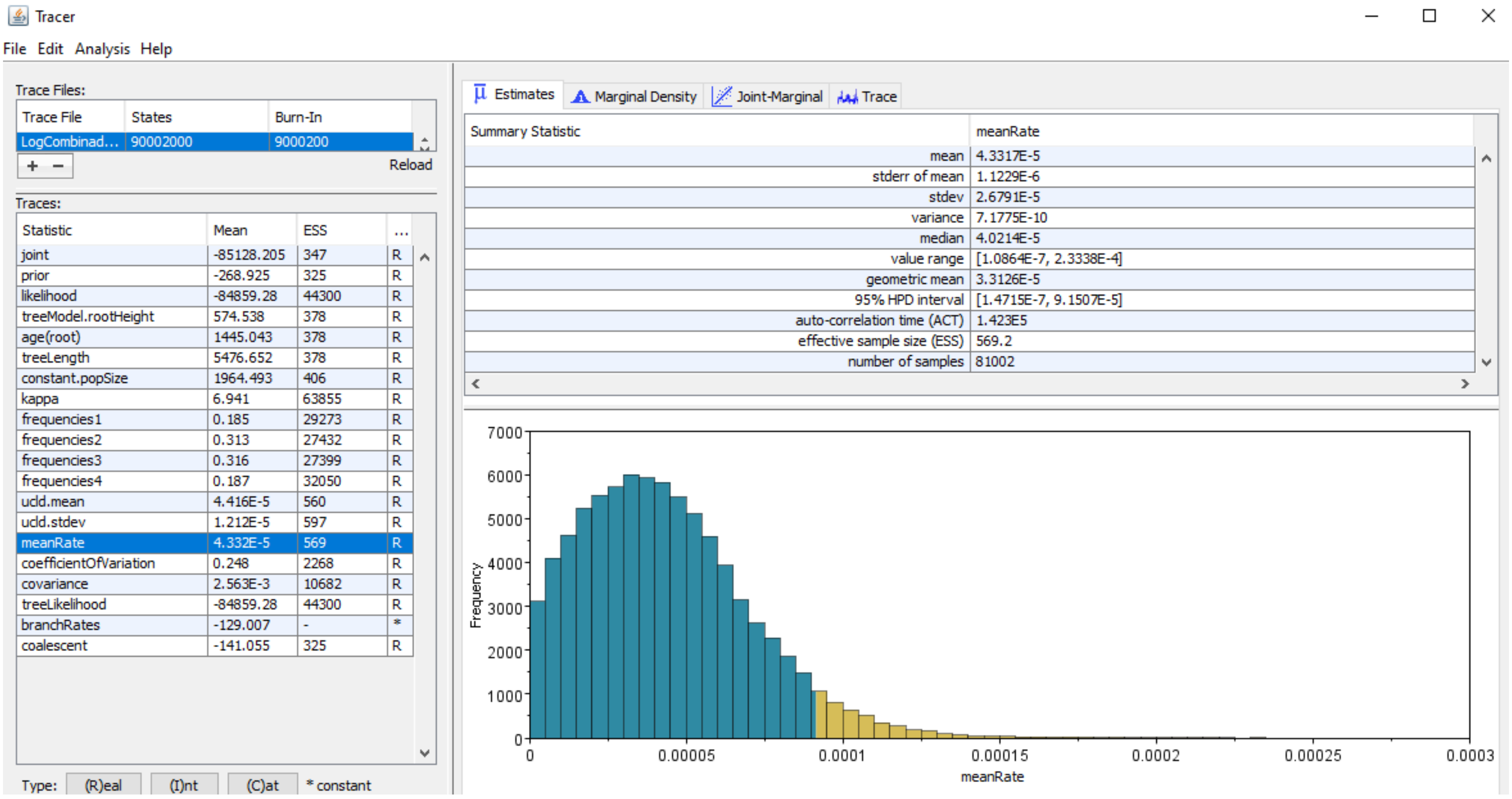


**MCC tree**

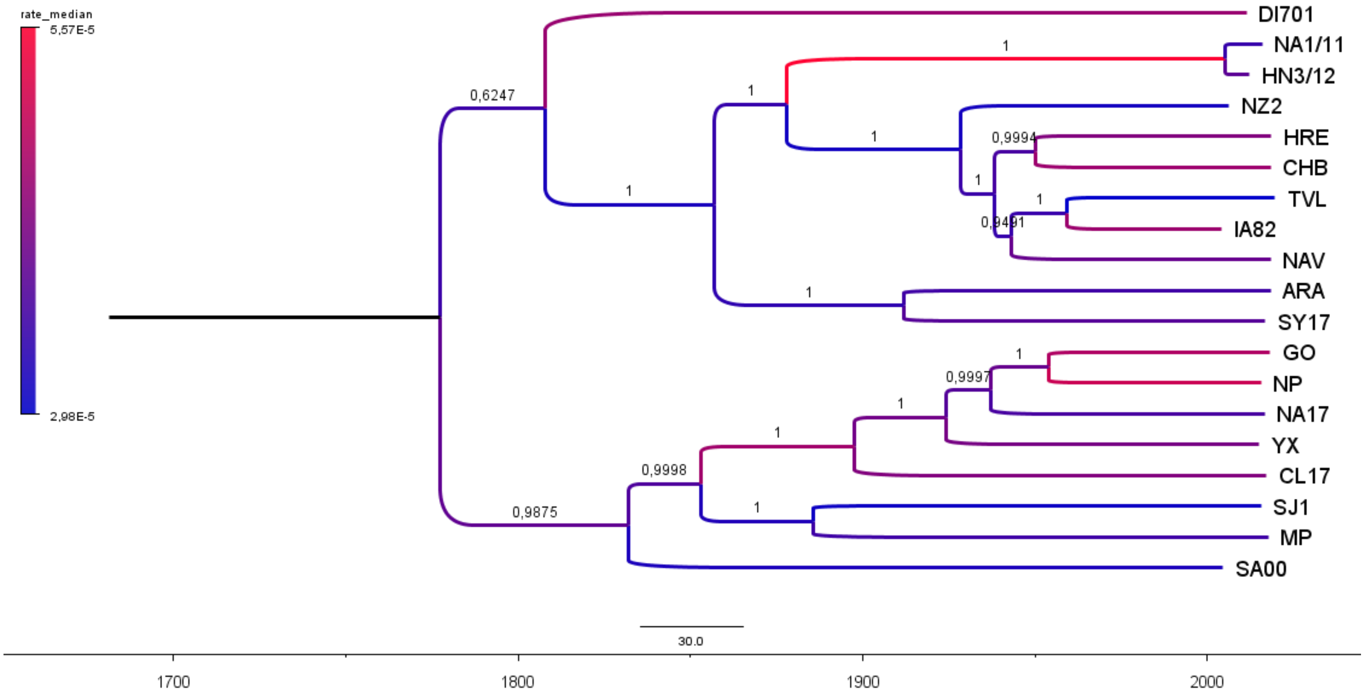

rate_median
5,57E-5
2,98E-5
DI701
NA1/11
HN3/12
NZ2
HRE
CHB
TVL
IA82
NAV
ARA
SY17
GO
NP
NA17
YX
CL17
SJ1
MP
SA00
0,6247
0,9875
0,9998
0,9994
0,9491
0,9997
30.0
1700
1800
1900
2000
